# Potential pitfalls of FISH microscopy as assessment method for anaerobic digesters

**DOI:** 10.1101/054999

**Authors:** Christian Abendroth, Sarah Hahnke, Michael Klocke, Olaf Luschnig

**Affiliations:** Robert Boyle Institute, Im Steinfeld 10, 07751 Jena, Germany; Bio H2 Umwelt GmbH, Im Steinfeld 10, 07751 Jena, Germany; Universidad de Valencia, Cavanilles Institute of Biodiversity and Evolutionary Biology, Spain; Leibniz Institute for Agricultural Engineering Potsdam-Bornim (ATB), Bioengineering, Max-Eyth-Allee 100, 14469 Potsdam, Germany

**Keywords:** FISH, Anaerobic Digestion, Biogas, Metabolic Activity, Microbial Activity, Fermenter diagnosis

## Abstract

In the present work we investigated how the state of a biogas reactor impacts the enumeration of prokaryotic cells by fluorescence *in situ* hybridisation (FISH). Therefore, the correlation between gas production and FISH hybridisation rates was analysed in different anaerobic digester sludges. High gasification activity coincided with high hybridisation rates. Low hybridisation rates were especially achieved with reactor samples subjected to long starvation periods showing low biogas production.

Based on our findings we conclude that samples for FISH analysis should be fixed as soon as possible to prevent a loss of microbial activity resulting in lower FISH signals. Furthermore, the location of sampling is of importance, since samples from different fermenters within the same biogas plant also varied strongly in their FISH hybridisation rate. Our results indicate that FISH could be a useful method for assessing the metabolic state of microorganisms in anaerobic digester plants.

## Introduction

The presented work is focused on the enumeration of bacteria and archaea in different anaerobic digester sludges using fluorescent *in situ* hybridisation (FISH). FISH is a well-established technique for targeting specific DNA or RNA sequences with oligonucleotide probes in microscopic samples. Publications dealing with FISH can be found back to the 70’s (e.g. in Cheung et al., 1977). Since this time the method has been improved and was used in many studies including the investigation of microbial communities of anaerobic biogas digesters. For example, FISH was used to elucidate the spatial organization of archaea and bacteria within methanogenic sludge granules in anaerobic reactors (Sekiguchi et al., 1999) and to analyse the methanogenic and bacterial population in a thermophilic anaerobic digester in response to changing concentrations of volatile fatty acids (Hori et al., 2006). The FISH approach was also used to quantify methanogenic archaea within different anaerobic digesters and was optimized in order to develop a protocol for flow-fluorescent *in situ* hybridisation, *i.e.* the combination of flow-cytometry and FISH (Nettmann et al., 2010 and 2013).

Based on this emerging interest in FISH as evaluation tool of anaerobic digestion processes, the aim of our study was to investigate whether FISH allows a reliable detection and thus a reliable enumeration of microorganisms in anaerobic digesters. Reviewing recent literature about the application of FISH in anaerobic digester sludges only poor information about this question was obtained. However, application of FISH in other fields of research resulted in findings that provide some hints concerning the efficiency of this technique. Manz et al. (1993) reported different FISH hybridisation rates for different samples of drinking water, which was linked to differences in the RNA content of the cells. It is commonly known that microbial activity is linked to the content of RNA. For example: A high growth rate of the bacterium *Vibrio natriegens* was linked to a high content of ribosomal RNA (Aiyar et al., 2002), furthermore it was shown that active, growing cells in the exponential growth phase contained stable ribosomes (Piir et al. 2011). On the other hand it is known that hampered growth conditions due to starvation lead to a decay of RNA (Deutscher, 2003; Zundel et al., 2009; Basturea et al., 2011; Piir et al., 2011). Thus, as FISH probes target rRNA molecules, it could be concluded that starvation might impair FISH hybridisation as well.

However, even though low RNA contents seem to be a good explanation for poor FISH enumeration results, there can be other reasons as well, such as difficulties in cell permeabilization due to differences in cell wall structures of gram positive and gram negative bacteria (Thurnheer et al., 2004) or signal falsification due to auto fluorescence from disturbing particles in the used sample (Bertaux et al., 2007).

In summary, based on previous publications we assumed that the enumeration of microbial cells in anaerobic digester plants by FISH might be insufficient depending on the reactor’s state and the metabolic state of the microorganisms, respectively. Such an effect would also be relevant when FISH is applied during long incubation periods of standardized gas production tests e.g. according to German VDI 4630 norm. To strengthen this hypothesis we performed a starvation experiment with reactor sludge to investigate whether reduced microbial metabolic activities result in fewer FISH signals. Furthermore, FISH hybridisation rates of different industrial reactor samples were analysed in order to evaluate the practical application of FISH-based evaluation analyses and whether FISH can be a useful tool to indicate reduced microbial activities in anaerobic digester plants.

## Methodology

### Sampling and analysis of biogas formation in different sludge samples

Reactor samples were taken from three different industrial anaerobic digesters (leach-bed systems). Leach-bed reactor 1 digested whole crop silage, corn silage, straw, shred, and cow manure. Leach-bed reactor 2 was supplied with municipal organic waste. Leach-bed reactor 3 was operated with corn silage, straw and cow manure. All biogas plants were described in detail previously by Abendroth et al. (2015). Additionally an agricultural two-stage system (one-phase), fed with silage, farm manure and livestock farming waste, especially designed to operate with high loading rates, was sampled. Samples were taken from the first stage (main digester designed as plug flow fermenter), from the second stage (completed stirred tank reactor, CSTR), and from the storage of digestion remnants.

Sludge was collected in 10 L plastic buckets for transportation and transferred immediately to the laboratory for analysis (Tab. S1). From all sludge samples the residual biogas production was analysed over a period of one week with an experimental setup according to the German guideline VDI 4630 (2006). Therefore, 0.5 L of the sludge samples was filled into 1 L batch-bottles directly upon sampling.

### Starvation experiment

A starvation experiment was set up with sludge from the two-stage biogas plant. Two 1 L bottles were filled with 0.5 L sludge each and incubated at 37°C. Formation of biogas was measured following the recommendations of the VDI 4630 (2006). After 18 days one of these assays (further referred to as control assay) was fed with 20 g of corn silage (fresh substrate). Afterwards, another starving period over 20 days followed. The second sample received no substrate and was subjected to starvation during the whole experiment. To analyse the effect of different temperatures on FISH hybridisation rates, one sludge sample was kept at room temperature (RT). Samples for FISH analyses were taken upon sampling of the biogas plant (day 0), before feeding (at day 17), upon feeding (at day 18), one day after feeding (at day 19) and after the second starvation period (at day 29).

The used corn silage was collected at the two-stage biogas plant and had 39.21% of total solids (TS) with 97.20% volatile solids (VS). TS and VS were analysed by gravimetric analysis. For determination of TS, samples were dried at 105°C. To determine VS, the samples were incinerated at 550°C (Nabertherm Model 4/11/R6, Germany).

### *Fluorescence* in situ *hybridisation (FISH)*

For FISH analyses 0.5 ml of the reactor samples were fixed with 1.5 ml 3.7% formaldehyde according to Nettmann et al. (2013). Samples of the leach-bed reactors and samples from the one-phase two-stage plant (reactor samples, which were compared to each other) were fixed for 8 h, due to the long transportation time of the sludge samples. Samples of the starvation experiment were fixed for 2 h. Incubation occurred in both cases at RT. After fixation samples were washed in 1x PBS (137 mM NaCl, 2.7 mM KCl, 10 mM Na_2_HPO_4_, 2 mM KH_2_PO_4_, pH 7.6) by centrifugation at 13,400 rpm for 10 min (Eppendorf Minispin, Germany) and stored in 48% ethanol (1x PBS and 96% ethanol mixed 1:1) at −20°C until further processing.

To perform FISH 100 μl of the samples were mixed with 500 μl 1x PBS and pelleted by centrifugation for 5 min at 15,000 rpm (Eppendorf Centrifuge 5417R), the supernatant was discarded and the pellet was resuspended in 1,000 μl 1x PBS. To support the dispersion of extracellular polymeric substances (EPS) a two-step ultrasonic treatment was carried out according to Nettmann et al. (2010). Afterwards the samples were washed in 1x PBS and resuspended in 100 μl of destilled water. Pre-treated samples were diluted (1:10, 1:50 or 1:100, depending on cell concentrations) and 10 μl of each sample were applied on teflon-coated slides covered with 0.1% gelatine and 0.01% CrK(SO_4_)_2_. Cells were permeabilized with lysozyme and achromopeptidase according to Sekar et al. (2003) with an extended lysozyme treatment of 90 min. Afterwards the slides were washed in destilled water, dried at 46°C, dehydrated in 96% ethanol.

For detection of bacteria a mix of the probes EUB338, EUB338II and EUB338III was used with 35% formamid in the hybridisation buffer (Daims et al., 1999; Glöckner et al., 1996), for detection of archaea the probe Arch915 was used (Stahl & Amann, 1991; Raskin et al., 1994) with 20% formamide. Nonspecific binding of labeled probes to the sample matrix or other nontarget objects was verified by using the nonsense probe NON338 (Wallner et al., 1993) under the same hybridisation conditions as the EUB338 probes. All probes were 5'– and 3'– end labeled with the fluorophore Cy3. Hybridisation was performed in a buffer-saturated humidity chamber according to Hugenholtz et al. (2002) with the following modifications: hybridisation at 46°C for 2 h with 50 ng probe, subsequent washing at 48°C for 20 min, washing buffer was supplemented with 5 mM EDTA. As additional control each hybridization was also performed with samples of pure cultures of a gram-positive bacterium (*Clostridium bornimense* strain M2/40^T^), a gram-negative bacterium (*Proteiniphilum* sp. strain M3/6) as well as an archaeal isolate (*Methanobacterium formicicum* strain L21–2). Counterstaining for total cell detection was performed with 4,6’–diamidino–2–phenylindole (DAPI) as described by Nettmann et al. (2010).

Samples were analyzed with a Nikon Optiphot–2 epifluorescence microscope (Nikon, Düsseldorf, Germany) equipped with the filter sets UV–2E/C and HQ:Cy3 for DAPI and Cy3 detection. For determination of total cell counts 12 view fields from independent, randomly chosen microscopic fields were counted at a magnification of 630x (approximately 1,500 cells).

### Calculation of hybridisation rates

For each sludge sample FISH and DAPI signals of 12 microscopic view fields for each, bacteria and archaea, were counted (in total 24 view fields). For each view field, the percentage of counted FISH signals to DAPI signals was calculated and a mean value for bacteria and archaea proportions was calculated for each sample. The percentage of the total amount of detected microorganisms was calculated by addition of the bacteria and archaea proportions (further just referred to as hybridisation rate). To calculate the standard deviation for the hybridisation rate, first the variance of SD_Bact_ and SD_Arch_ was calculated, afterwards the square root of the summed up variances was formed.

## Results and discussion

### Impact of starvation on FISH hybridisation rates

To investigate how microbial activities impact FISH hybridisation rates (*i.e.* the proportion of microbial cells detected by FISH), a starvation experiment was performed. Therefore, sludge from a mesophilic biogas plant was incubated for 29 days at 37°C without feeding and FISH hybridisation rates were determined at different time points. As a control a parallel assay was fed with fresh corn silage at day 18. During the first starvation period (up to day 18) both assays exhibited a similar biogas production increasing up to 11,450 mL L^−1^ and 13,130 mL L^−1^, respectively. The minor difference in produced biogas might be explained due to high inhomogeneity of the used sludge sample. Feeding of the control assay at day 18 led to an abrupt increase in biogas production reflecting stimulation of microbial activity, whereas the untreated sample showed a continuing small increase in biogas formation (Fig. 1A).

**Figure 1.**
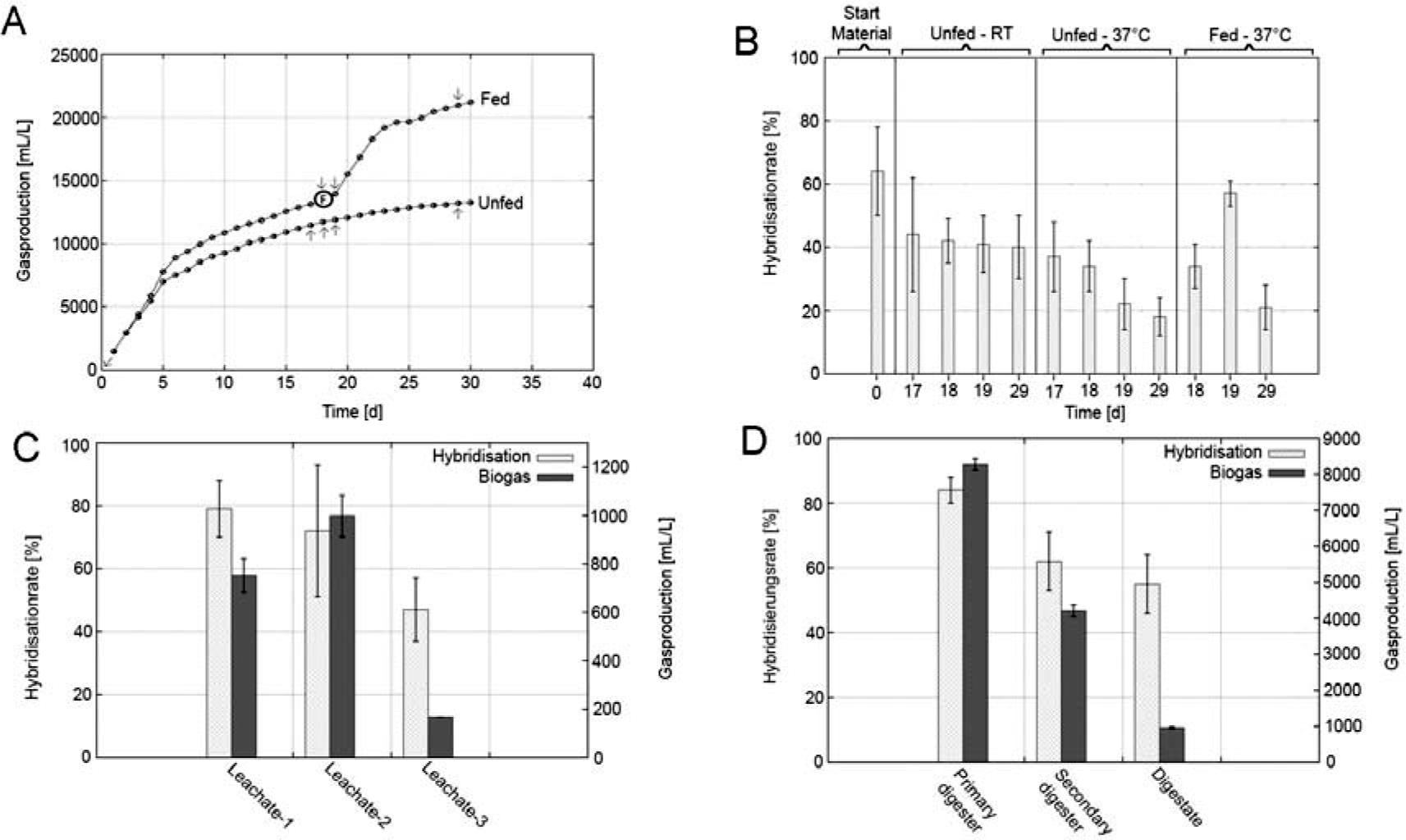
*Impact of sludge activity on FISH hybridisation. (A) Sampling overview and biogas production during the starvation experiment incubated at 37°C with sludge from the main digester of the two-stage system. “F” indicates feeding of the control assay with 20 g fresh corn silage, arrows indicate sampling points for FISH analysis. (B) FISH hybridisation rates (proportions of FISH signals to DAPI counts) in sludge samples subjected to starvation (incubated at RT and at 37°C) and in the control assay fed with fresh substrate at day 18 (incubated at 37°C). (C) Produced volumes of biogas and FISH hybridisation rates of* leachate from three different industrial leach-bed systems. Values of produced biogas volumes originate from Abendroth et al. (2015). (D) Produced volumes of biogas and FISH hybridisation rates of three different stages of a two-stage reactor system.

The highest FISH hybridisation rate of 64% was achieved at the start of the experiment with sludge directly taken from the biogas plant. Elongated incubation of the sludge under starvation conditions led to a significant decrease in FISH hybridisation rates reaching 37.5% at day 17 in the sample incubated at 37°C, and 44.2% in a control assay incubated at room temperature (RT), respectively (Fig. 1B, Tab. 1).

Directly upon feeding at day 18 FISH analyses of the fed and the unfed assay (at 37°C) resulted in similar hybridisation rates showing no significant differences (Tab. 1). However, after one day of incubation with fresh substrate the hybridisation rate increased significantly from 35.0% to 57.8%, a proportion similar to the initial hybridisation rate (at day 0) obtained with the reactor material directly taken from the biogas plant. In contrast, the hybridisation rate in the assay subjected to further starvation continued to decrease over time (Fig. 1B). Eleven days after feeding the hybridisation rate of the fed sample was again low coinciding with a declined biogas formation indicating low microbial activity. With 21.5% the hybridisation rate was comparable to that obtained in the untreated sample (18.1%). FISH hybridisation rates of the unfed assay incubated at 37°C decreased significantly faster than that of the unfed control at RT. It is known that incubation at room temperature can lower the biogas production (Prasad et al. 2012). Thus, it could be assumed that the microorganisms at 37°C exhibited higher metabolic activities resulting in a faster turnover of the remaining substrates and nutrients compared to those incubated at RT.

**Table 1.**
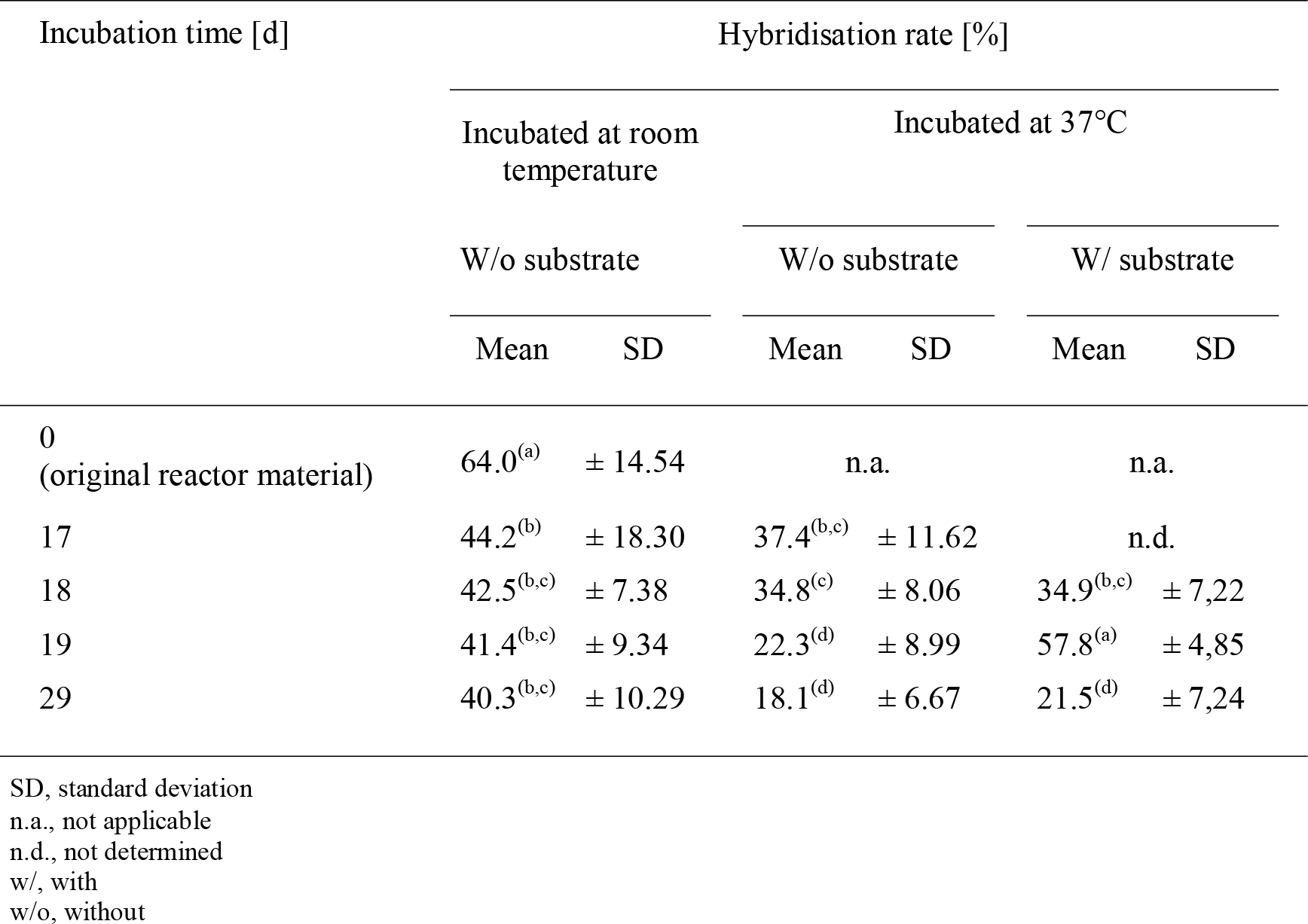
*FISH hybridisation rates with respect to the duration of starvation. The letters in parenthesis show the results of statistical analysis (Duncan’s multiple range test, p<0.05); different letters indicate significant difference among the samples.*

### Comparison of industrial digesters

As the lab-scale experiment with digester sludge revealed a negative effect of long starvation periods on FISH hybridisation rates, we performed FISH with reactor samples of different industrial biogas reactors to investigate, whether a correlation between reactor activity and FISH hybridisation efficiencies can also be observed with samples directly taken from large-scale biogas plants. Leachates from three different leach-bed systems were compared with respect to the produced biogas amounts and the corresponding FISH hybridisation rates. The samples from the leach-bed reactor fed with whole crop silage, corn silage, straw, shred, and cow manure (Leachate–1) and the reactor fed with municipal organic waste (Leachate–2) showed a much higher gas production compared to the leachate from the reactor fed with corn silage, straw, and cow manure (Leachate–3); namely 751 mL and 998 mL compared to 166 mL biogas per litre of sludge sample. The differences in the produced biogas volumes can be explained by differences in the proportions of biologically degradable organic matter in the supplied substrates. Chemical analyses revealed, that Leachate–1 and Leachate–2 contained higher concentrations of organic acids (747 mg L^−1^ and 1,170 mg L^−1^ total volatile fatty acids [TVFA]), which can be converted into biogas, than Leachate–-3, in which no TVFA could be detected (Tab. S1). This can be explained due to the fact that the leach-bed system from Leachate–3 was mainly fed with cow manure (up to 70%), while the other leach-bed systems contained high amounts of energy-rich substrates like corn silage and domestic waste. In accordance with the assumption that high gas production is linked to an active metabolic state of the microorganisms, samples with high gas volumes coincided with high FISH hybridisation rates. FISH hybridisation rates of 79% and 72% were obtained with the samples Leachate–1 and Leachate–2 whereas in Leachate–3 only 48% of the total DAPI counts could be detected by FISH (Fig. 1C, Tab. 2).

**Table 2.** *Statistical comparison between FISH hybridisation rates of samples from different industrial leach-bed biogas reactors and of samples from different stages of a two-stage reactor system. The letters in parenthesis show the results of statistical analysis (Duncan’s multiple range test, p<0.05); different letters indicate significant difference among the samples.*

| Sampled leach-bed reactors | Mean | SD | Sampled stages of the two-stage reactor system | Mean | SD |
| --- | --- | --- | --- | --- | --- |
| Leachate-1 | 79.1<sup>(a,b)</sup> | ±9.67 | Primary Digester | 84.3<sup>(b)</sup> | ±4.95 |
| Leachate-2 | 72.2<sup>(a)</sup> | ±21.45 | Secondary Digester | 62.1<sup>(d)</sup> | ±9.85 |
| Leachate-3 | 47.9<sup>(c)</sup> | ±1.93 | Digestate | 55.1<sup>(c,d)</sup> | ±9.07 |

SD, standard deviation

Comparing sludges from different stages from a one-phase, two-stage plant (primary digester, secondary digester, and digestate) a similar correlation between the biogas production and achieved FISH hybridisation rates was observed: high FISH hybridisation rates were achieved with sludges showing high biogas production. The primary digester showed the highest gas production (8,270 ml L^−1^) and a FISH hybridisation rate of 84% was achieved. In the secondary digester a lower gas production was measured (4,207 ml L^−1^) and the obtained FISH signals accounted for 62% of the total DAPI cell counts. The lowest biogas production was measured in the digestate (955 ml L^−1^) in which the FISH hybridisation rate showed only 55% (Fig. 1D). Again the observed volumes of produced biogas are in accordance with environmental chemical parameters, since samples with higher volumes of produced gas also contained more TVFA (Tab. S1).

All sampled reactors showed an optimal range of pH and no accumulation of inhibitory organics acids (Table S1). Predominance of acetic acid was observed. Samples with high volumes of produced biogas also showed high concentrations of organic acids. These parameters indicate that the sludge samples were in good and undisturbed conditions. There is a low probability that inhibitory effects (other then low amounts of nutrients) might lower metabolic activity and therefore lower hybridisation rate.

Our results demonstrate that FISH hybridisation rates can differ substantially between different sludge samples. The fact that samples with highest gas production also showed highest FISH hybridisation rates indicates a relation between the microbial metabolic activity and achieved FISH signals. However, as the hybridisation rates in our study never exceeded 80% even in the sludges that produced the highest amounts of biogas (Leachate–1, Leachate–2 and primary digester), it can be suggests that other reasons than metabolic activity influenced the hybridisation efficiency as well (e.g. organism-specific permeabilization of cell walls).

## Conclusions

Our results show that FISH hybridisation rates, i.e. the percentage of microorganisms detected by FISH, can differ strongly between sludges of different anaerobic digesters and hybridisation rates were strongly influences by the metabolic state of the microorganisms. Thus, applying FISH as enumeration tool for different groups of microorganisms in anaerobic digester sludges, it must be considered that cell numbers can be error prone as the result of insufficient microbial metabolic activity. Especially long starvation periods of sludge samples reduced the amount of detectable microorganisms, which must also be taken into consideration when FISH is used in combination with other methods for an overall reactor screening such as a gas production assay, e.g. according to German VDI 4630 guidelines. Long incubation and periods during the gas measurement would probably lead to microbial starvation and thus would falsify FISH results. In general, in case of application of FISH as screening tool we recommend to select samples from most active reactor phases, since less active phases like the digestate storage showed lower hybridisation rates. Furthermore, our results indicate that FISH could be used to assess the metabolic state of microorganisms in anaerobic digester plants. In case of low microbial activity the reactor performance could be improved by addition of substrate.

## Acknowledgements

We thank Claudia Semionov and Robert Förster for technical support. This work was supported by a grant of the German Federal Ministry of Economics and Technology [ZIM, grant 16KN017626 and 16KN017629].

